# All-polymeric transient neural probe for prolonged in-vivo electrophysiological recordings

**DOI:** 10.1101/2021.03.09.434622

**Authors:** Laura Ferlauto, Paola Vagni, Elodie Geneviève Zollinger, Adele Fanelli, Katia Monsorno, Rosa Chiara Paolicelli, Diego Ghezzi

## Abstract

Transient bioelectronics has grown fast, opening possibilities never thought before. In medicine, transient implantable devices are interesting because they could eliminate the risks related to surgical retrieval and reduce the chronic foreign body reaction. However, despite recent progress in this area, the short functional lifetime of devices due to short-lived transient metals, which is typically a few days or weeks, still limits the potential of transient medical devices. We report that a switch from transient metals to an entirely polymer-based approach allows for a slower degradation process and a longer lifetime of the transient probe, thus opening new possibilities for transient medical devices. As a proof-of-concept, we fabricated all-polymeric transient neural probes that can monitor brain activity in mice for a few months rather than a few days or weeks. Also, we extensively evaluated the foreign body reaction around the implant during the probe’s degradation. This kind of devices might pave the way for several applications in neuroprosthetics.

## Introduction

Building devices able to disappear in the surrounding environment after a programmed lifetime, leaving minimal and harmless traces after their disposal, is a fascinating idea for the development of green electronics to preserve the environment by reducing waste production and recycling processes[1–5]. The same concept is also appealing for medical devices[6–8], such as localized drug release systems[9], power units[10], external sensors[11], and implantable recording systems[12] to eliminate the risks related to surgical retrieval[13] and reduce the chronic foreign body reaction[14].

In medical applications, the durability of transient devices is highly dependent on the degradation time and process of the materials used for the substrate and encapsulation layers and, most importantly, of the functional components exposed to the body. While several biodegradable natural or synthetic polymers, such as cellulose[1], silk[15] and poly(lactic-co-glycolic acid)[16,17], are often used as substrate and encapsulation materials, the choice for the functional elements has always landed so far on transient inorganic materials. These materials, such as magnesium, zinc, molybdenum, or iron, dissolve within minutes, hours, or maximum days, once in contact with body fluids[11,18]. This short functional window still limits the list of potential medical applications for transient devices.

To extend the lifetime of transient medical devices, and consequently increase their possible applications, such as for example mid-term monitoring of the brain activity, our strategy is to switch from transient metals to entirely polymer-based devices. Our results show the fabrication and the in-vitro characterization of all-polymeric transient neural probes (TNPs) allowing prolonged electrophysiological recordings in-vivo up to three months post implantation. Histological studies revealed minimal foreign body reaction both in short-term and long-term.

## Materials and Methods

### Fabrication of transient all-polymeric neural probe

Two 4-inch silicon wafers (thickness 525 μm) were etched (AMS 200, Alcatel) to obtain a protruding wafer including electrodes, traces, and pads, and a hollow wafer with only traces (both 100 μm depth). Polycaprolactone (MW 50,000; 25090, Polyscience, Inc.,) in powder was dissolved in chloroform (C2432, Sigma-Aldrich) at a concentration of 0.25 mg ml^-1^ and was left stirring and heating at 60 °C overnight over a hotplate. The day after, 6 ml of polycaprolactone solution were spin-coated on both silicon moulds (protruding: 150 rpm for 30 sec; hollow: 200 rpm for 30 sec) pre-treated with chlorotrimethylsilane (92361, Sigma-Aldrich) to prevent permanent attachment of the polymer layer. The wafers were then left 30 min in the oven at 75 °C to let the chloroform evaporate. After a cool-down period of at least 2 hours, the polycaprolactone layer from the protruding wafer was peeled off, and the design (electrodes, traces and pads) was filled with an aqueous solution of PEDOT:PSS (M122 PH1000, Ossila) mixed with 20 wt% ethylene glycol (324558, Sigma-Aldrich) and 1 wt% (3-Glycidyloxypropyl)trimethoxylane (440167, Sigma-Aldrich) before curing overnight in an oven at 37 °C. The polycaprolactone layer from the hollow wafer, instead, was peeled off after laser-cutting the openings for the electrodes and pads (Optec MM200-USP). Eventually, the two layers were manually aligned and fused together over a hotplate at 60 °C for a few seconds. A silver-based epoxy (H27D Kit Part A, Epo-Tek) was then applied on the pads and connectors were positioned and immobilized with silicone (DC 734 RTV clear, Dow Corning). Three types of all-polymeric transient neural probes were fabricated with a different number of electrodes: 17 electrodes (700 μm in diameter; connector 61001821821, Wurth Elektronik), 4 electrodes (500 μm in diameter; connector 143-56-801, Distrelec), and 1 electrode (500 μm in diameter; no connector). Samples with fluorescent PEDOT:PSS were prepared as previously described[19]. Briefly, poly(vinyl alcohol) (MW 130,000; 563900, Sigma-Aldrich) was dissolved in deionization water 5 wt%; a composite solution (25 wt%) of poly(vinyl alcohol) and PEDOT:PSS:EG was prepared, deposited on the polycaprolactone, and baked (50 °C, 10 min). (3-Glycidyloxypropyl)trimethoxylane (440167, Sigma-Aldrich) was then deposited by vapour deposition at 90 °C for 45 min. Afterwards, fluorescein isothiocyanate labelled poly-L-lysine (50 μg ml^-1^ in phosphate-buffered saline; P3543, Sigma-Aldrich) was drop-casted on the sample and allowed to react for 2 hr at room temperature. The sample was then rinsed with phosphate buffered saline (0.1 M), sodium chloride (0.1 M), and deionized water.

### Electrochemistry

Electrochemical characterization was performed with a three-electrode potentiostat (Compact Stat, Ivium) at room temperature. Each all-polymeric transient neural probe was immersed in phosphate-buffered saline (pH 7.4) together with a silver / silver chloride reference electrode and a platinum counter electrode. Impedance spectroscopy was measured between 1 Hz and 1 MHz using an AC voltage of 50 mV. With the same setup, cyclic voltammetry was obtained by sweeping a cyclic potential at a speed of 50 mV s^-1^ between −0.9 and 0.8 V. For each electrode, the average response over five cycles was calculated, and the total charge storage capacity was computed from the integration of the respective currents.

### Scanning electron microscopy

Images of the topography and cross-section of the all-polymeric transient neural probes were taken with a Schottky field emission scanning electrons microscope (SU5000, Hitachi) and image post-processing was performed with ImageJ.

### Cytotoxicity test

A test on extract was performed on two all-polymeric transient neural probes (4-electrodes design) sterilized by UV exposure, with a ratio of the product to extraction vehicle of 3 cm^2^ ml^-1^. The extraction vehicle was Eagle’s minimum essential medium (Thermo Fisher Scientific, 11090081) supplemented with 10% fetal bovine serum (Thermo Fisher Scientific, 10270106), 1 % penicillin-streptomycin (Thermo Fisher Scientific, 15070063), 2 mM L-Glutamine (Thermo Fisher Scientific, 25030081), and 2.50 μg ml^-1^ Amphotericin B (Gibco-Thermo Fisher Scientific, 15290026). The extraction was performed for 24 hr at 37°C and 5% CO_2_. L929 cells (Sigma, 88102702) were plated in a 96 well plate at a sub-confluent density of 7000 cells per well in 100 μl of the same medium. L929 cells were incubated for 24 hr at 37°C and 5% CO_2_. After incubation, the medium was removed from the cells and replaced with the extract (100 μL per well). After another incubation of 24 hr, 50 μL per well of XTT reagent (Cell proliferation kit 11465015001, Sigma) were added and incubated for 4 hr at 37 °C and 5% CO_2_. An aliquot of 100 μl was then transferred from each well into the corresponding wells of a new plate, and the optical density was measured at 450 nm by using a plate reader (FlexStation3, MolecularDevices). Clean medium alone was used as a negative control, whereas medium supplemented with 15% of dimethyl sulfoxide (D2650-5X5ML, Sigma) was used as a positive control. Each condition was tested in triplicates.

### Accelerated degradation

All-polymeric transient neural probes were prepared as described above, with the 4-electrode design. After fabrication, samples were weighted to set the initial weight value before starting degradation. Degradation of each sample occurred in 10-ml phosphate-buffered saline at 37 °C and was accelerated by pH increase: 100μl of NaOH (2M NaOH Standard solution, 71474-1L, Fluka) were added to the solution to reach pH 12. At each time point, samples were first gently dried with a tissue and then let dry completely under vacuum at room temperature for 4 hr. Once dry, samples were weighted. Normalized weight for each sample was computed as the ratio between the weight at each time point and the initial value.

### Animal handling

All experiments were conducted according to the animal authorizations GE13416 approved by the Département de l’emploi, des affaires sociales et de la santé (DEAS), Direction générale de la santé of the Republique et Canton de Geneve in Switzerland and VD3420 approved by Service de la consommation et des Affaires vétérinaires (SCAV) of the Canton de Vaud in Switzerland. All the experiments were carried out during the day cycle. For the entire duration of the experiment, the health condition was evaluated three times a week, and the bodyweight was controlled once a week. Experiments were performed in adult (> 1-month-old) mice. Male and female C57BL/6J mice (Charles River) were kept in a 12 h day/night cycle with access to food and water *ad libitum.* White light (300 ± 50 lux) was present from 7 AM to 7 PM and red light (650-720 nm, 80-100 lux) from 7 PM to 7 AM. Homozygous B6.129P2(Cg)-Cx3cr1^tm2.1(cre/ERT2)Litt^/WganJ mice (The Jackson Laboratory, Stock No: 021160) and homozygous B6.Cg-Gt(ROSA)26Sor^tm14(CAG-tdTomato)Hze^/J mice (The Jackson Laboratory, Stock No: 007914) were kept in a 12 h day/night cycle with access to food and water *ad libitum.* White light (300 ± 50 lux) was present from 7 AM to 7 PM and no light from 7 PM to 7 AM. Homozygous B6.129P2(Cg)-Cx3cr1^tm2.1(cre/ERT2)Litt^/WganJ mice and homozygous B6.Cg-Gt(ROSA)26Sor^tm14(CAG-tdTomato)Hze^/J mice were crossed to obtain heterozygous mice bearing a tamoxifen-inducible expression of the tandem dimer Tomato fluorescent in microglia in the brain. Mice were injected with tamoxifen dissolved in corn oil (75 mg kg^-1^) to induce the expression at postnatal day 35. Surgery was performed between 2 weeks and 1 month after.

### Surgical implantation

Mice were anesthetized with isoflurane inhalation (induction 0.8-1.5 l min^-1^, 4-5%; maintenance 0.8-1.5 l min^-1^, 1-2%). Analgesia was performed by subcutaneous injection of buprenorphine (Temgesic, 0.1 mg kg^-1^), and a local subcutaneous injection of lidocaine (6 mg kg^-1^) and bupivacaine (2.5 mg kg^-1^) with a 1:1 ratio. The depth of anaesthesia was assessed with the pedal reflex, and artificial tears were used to prevent the eyes from drying. The temperature was maintained at 37 °C with a heating pad during both surgical procedures and recording sessions. The skin of the head was shaved and cleaned with betadine. Mice were then placed on a stereotaxic frame, and the skin was opened to expose the skull. A squared craniotomy of approximately (4 x 4 mm) was opened over the visual cortex (identified by stereotaxic coordinates), and the dura mater was removed. UV-sterilized transient all-polymeric neural probes were inserted in the cortex using a micromanipulator (SM-15R, Narishige). The probes were always implanted with the PEDOT:PSS:EG layer exposed to the caudal part of the brain to clearly discriminate the two sides during image analysis. A reference screw electrode was implanted in the rostral side of the cranium. The craniotomy was closed using dental cement, and an extra layer of dental cement was applied to secure the probe. The mice were left to recover on a heating pad and later returned to their cage.

### Induction of seizure and monitoring of epileptic activity

Immediately after surgery, mice were removed from the stereotaxic apparatus while still under general anaesthesia, a needle electrode was placed subcutaneously in the dorsal area near the tail as ground, and the four channels of transient all-polymeric neural probes and the reference electrode were connected to the amplifier (BM623, Biomedica Mangoni). The signals were recorded using the WinAver software (Biomedica Mangoni). The recording was started (bandpass filtered 0.1-2000 Hz and sampling frequency 8192 Hz) and, after a baseline period, 20 μl of pentylenetetrazol (50 mg ml^-1^, 45 mg kg^-1^) were delivered via intraperitoneal injection. Recordings were maintained during the ictal events. The animals were euthanized with an injection of pentobarbital (150 mg kg^-1^).

### Chronic recordings

A first acute recording session was carried out immediately after the implantation, in order to check the functionality of the probes at the starting point of the experiment. Chronic recordings were then performed 1- and 2-weeks post-implantation and then every 2 weeks until 12 weeks. The mice were dark-adapted for two hours before each chronic recording session. Mice were anaesthetized with a mixture of ketamine (87.5 mg kg^-1^) and xylazine (12.5 mg kg^-1^). The depth of anaesthesia was assessed with the pedal reflex, and artificial tears were used to prevent the eyes from drying. The temperature was maintained at 37 °C with a heating pad during both surgical procedures and recording sessions. Mice were placed on a stereotaxic apparatus in front of a Ganzfeld flash stimulator (BM6007IL, Biomedica Mangoni) and a needle electrode was placed subcutaneously in the dorsal area near the tail as ground. The four channels of transient all-polymeric neural probes and the reference electrode were connected to the amplifier (BM623, Biomedica Mangoni). The signals were recorded using the WinAver software (Biomedica Mangoni). First, a baseline recording was obtained for 2.5 min (bandpass filter 0.1 - 2000 Hz and sampling frequency 8192 Hz). The recording noise was extracted from the baseline recording using a moving average window of 10 points; the mean ± s.d. was calculated for each 500 ms epoch, then the mean s.d. of all the epochs was averaged to obtain the noise of the whole recording. Channels with a noise level higher than the average plus twice the s.d. of the noise at each time point were considered as non-working channels and excluded from further analysis. Local field potentials were then acquired for 2.5 min (bandpass filter 0.1 - 100 Hz and sampling frequency 819.2 Hz). After the recordings, a high pass filter at 0.5 Hz was applied, and the periodogram was calculated using the Welch method (window of 4 s) on the whole recording. The area below the curve between 0.5 and 4 Hz frequencies (corresponding to the delta band) was approximated using the composite Simpson’s rule. The power of the delta band was then divided by the total power (area below the periodogram) to obtain the relative power of the delta band. Last, visually evoked potentials were recorded (bandpass filter 0.1-200 Hz and sampling frequency 2048 Hz) upon the presentation of 10 consecutive flashes (4 ms, 10 cd s m^-2^) delivered at 1 Hz of repetition rate. Data were processed and analysed using MATLAB (MathWorks). After each session, the mice were left to recover on a heating pad and later returned to their cage.

### Histological analysis

Animals were euthanized with an injection of pentobarbital (150 mg kg^-1^) under a chemical hood. The chest cavity was opened to expose the beating heart, and a needle was inserted in the left ventricle, while the right atrium was cut to allow complete bleeding. The animal was immediately perfused with phosphate-buffered saline followed by a fixative solution of 4% paraformaldehyde in phosphate buffered solution. At the end of the procedure, the head of the animal was removed, and the brain collected and placed in 4% paraformaldehyde overnight for post-fixation. For section preparation, the samples were cryoprotected in sucrose 30 % and frozen in OCT. 20-μm thick horizontal sections of the brain were obtained using a cryostat (Histocom, Zug, Switzerland) and placed on microscope slides. The sections were washed in phosphate-buffered saline, permeabilized with phosphate-buffered saline + Triton 0.1 % (Sigma-Aldrich), left for one hour at room temperature in blocking buffer (Triton 0.1 % + 5 % normal goat serum), and incubated overnight at 4 °C with primary antibodies: Anti-GFAP 1:1000 (Z0334, Dako), Anti-CD68 1:400 (MCA1957, Biorad), and Anti-NeuN 1:500 (ABN90P, Millipore). The day after, the sections were incubated for two hours at room temperature with secondary antibodies 1:500 (Alexa Fluor 647 and 488, Abcam), counterstained with DAPI 1:300 (Sigma-Aldrich), and mounted for imaging with Fluoromount (Sigma-Aldrich) solution. Image acquisition was performed with a confocal microscope (LSM-880, Zeiss). For image segmentation and quantification, images were acquired using a slide scanner microscope (Olympus VS120) and analysed in Python, using the scikit-image package (https://scikit-image.org/). First images were rotated to align the probe horizontally. Each image was converted to binary using a local threshold algorithm: all pixels whose intensity was above the threshold were assigned the value 1, while to the rest of the pixels it was assigned the value 0. A group of adjacent pixels with value 1 was defined as a blob, and the area of each blob was calculated, along with the coordinates of its centroid. The image was divided into 10 x 10 quadrants, for each of which the total area of blobs (with their centroid included in that quadrant) was computed. Then, the total area occupied by blobs in the zone adjacent to the probe (proximal area) was calculated as follows. First, the proximal area was calculated as the cumulative area of all the quadrants included in a rectangle defined by the extremities of the probe (manually specified for each of the images, see **Supplementary Fig. 4**), minus the area of the probe itself. From this, the percentage of this area occupied by blobs was computed as the area occupied by blobs over the proximal area. Finally, the average percentage of the proximal area occupied by the blobs for each sample was used as a parameter to compare the experimental groups. For the analysis of CD68 signal at the electrode level (**Supplementary Fig. 5**), each image was divided into 10 x 10 quadrants, as described above, and for each of them, the total area of blobs (with their centroid included in that quadrant) was computed. Then, the image was divided into two parts: the side of the probe with the PEDOT:PSS:EG exposed and the side of the probe with only the PCL exposed. For each side, the cumulative area occupied by blobs was calculated along the x-axis and normalized by the global maximum.

### Whole-brain imaging

Brain clarification was performed according to a previously described procedure[20]. Briefly, after overnight post-fixation at 4 °C in PFA 4%, the brain was immersed in hydrogel solution (Acrylamide 40% + VA-044 initiator powder in phosphate-buffered saline) at 4 °C for three days. The hydrogel polymerization was induced by keeping the sample at 37 °C for 3 hr. Then, the brain was passively clarified in 4% sodium dodecyl sulfate clearing solution (pH 8.5) for four weeks under gentle agitation at 37 °C. The whole clarified brain was transferred to Histodenz solution at pH 7.5 (Sigma). Brains were immersed in a refractive index matching solution (RIMS) containing Histodenz for at least 24 hours before being imaged. Brains were glued to a holder and immersed in a 10 x 20 x 45 mm quartz cuvette filled with RIMS. The cuvette was then placed in a chamber filled with oil with (n = 1.45; Cargille). A custom-made light-sheet microscope optimized for labelled clarified tissue was used (Clarity Optimized Light-sheet Microscope[21]) to image the implant within the mouse brain (**Fig. 6b,c**). The sample was illuminated (488 and 554 nm) by two digitally scanned light-sheets coming from opposite directions, and the emitted fluorescence was collected by high numerical aperture objectives (Olympus XLPLN10XSVMP, N.A 0.6) filtered (Brightline HC 525/50, Semrock) and imaged on a digital CMOS camera (Orca-Flash 4.0 LT, Hamamatsu) at a frequency ranging between 5 and 10 frames per second. A self-adaptive positioning of the light sheets across Z-stacks acquisition ensured an optimal image quality over up to 1 cm of tissue. To acquire images of whole samples at lower resolution (**Fig. 6a**) anther custom-made light-sheet microscope was used (mesoscale selective plane illumination microscopy[22]). The microscope consists of a dual-sided excitation path using a fibre-coupled multiline laser combiner (405, 488, 561 and 647 nm; Toptica MLE) and a detection path comprising a 42 Olympus MVX-10 zoom macroscope with a 1x objective (Olympus MVPLAPO), a filter wheel (Ludl 96A350), and a CMOS camera (Hamamatsu Orca Flash 4.0 V3). The excitation paths also contain galvo scanners for light-sheet generation and reduction of shadow artefacts due to absorption of the light-sheet. In addition, the beam waist is scanned using electrically tuneable lenses (ETL, Optotune EL-16-40-5D-TC-L) synchronized with the rolling shutter of the CMOS camera. This axially scanned light-sheet mode (ASLM) leads to a uniform axial resolution across the field-of-view of 5 μm. Image acquisition was made using custom software written in Python. Z-stacks were acquired at 3 μm spacing with a zoom set at 2x resulting in an inplane pixel size of 7.8 μm (2048×2048 pixels). The excitation wavelength was set at 561 nm with an emission 530/40 nm bandpass filter (BrightLine HC, AHF).

### Statistical analysis and graphical representation

Statistical analysis and graphical representation were performed with OriginPro 2017 (OriginLab) and Prism 8 (Graph Pad). The normality test (D’Agostino & Pearson omnibus normality test) was performed in each dataset to justify the use of a parametric or nonparametric test. In each figure p-values were represented as: * p < 0.05, ** p < 0.01, *** p < 0.001, and **** p < 0.0001.

### Data availability

The authors declare that all other relevant data supporting the findings of the study are available in this article and in its supplementary information file. Access to our raw data can be obtained from the corresponding author upon reasonable request.

## Results

### Probe fabrication and characterisation

As a proof-of-concept, we fabricated TNPs based on polycaprolactone (PCL) as substrate and encapsulation layers, and poly(3,4-ethylenedioxythiophene)-poly(styrenesulfonate) (PEDOT:PSS) doped with ethylene glycol (EG) as a conductive element, by using soft lithographic techniques (**Fig. 1a,b** and **Supplementary Fig. 1**). PCL is a commercial biodegradable polyester widely used in biomedical applications[23–26]. It is easy to process and has a longer degradation time (several months or years[23]) compared to most biodegradable polymers. PEDOT:PSS dispersed in water is commercially available, has excellent biocompatibility, has the ability to support cells viability[27,28], and can be integrated into softer systems like elastomers[29,30] and hydrogels[31,32]. EG doping of PEDOT:PSS is known to improve its conductivity[33,34]. The micro-structured PCL-based substrate (thickness 60 to 70 μm) and encapsulation (thickness 60 to 70 μm) layers were obtained by replica moulding. The hollow regions of the substrate layer were then manually filled with PEDOT:PSS doped with EG (PEDOT:PSS:EG) to create electrodes, traces, and contact pads (**Supplementary Fig. 1**). At the electrode level, the PEDOT:PSS:EG layer is approximately 3-μm thick, and at the trace level, it is approximately 1-μm thick (**Supplementary Fig. 2**). After curing of the PEDOT:PSS:EG, the PCL encapsulation layer was manually flip-bonded to the substrate layer via a low-temperature process (total probe thickness ranging from 120 to 140 μm).

**Figure 1.**
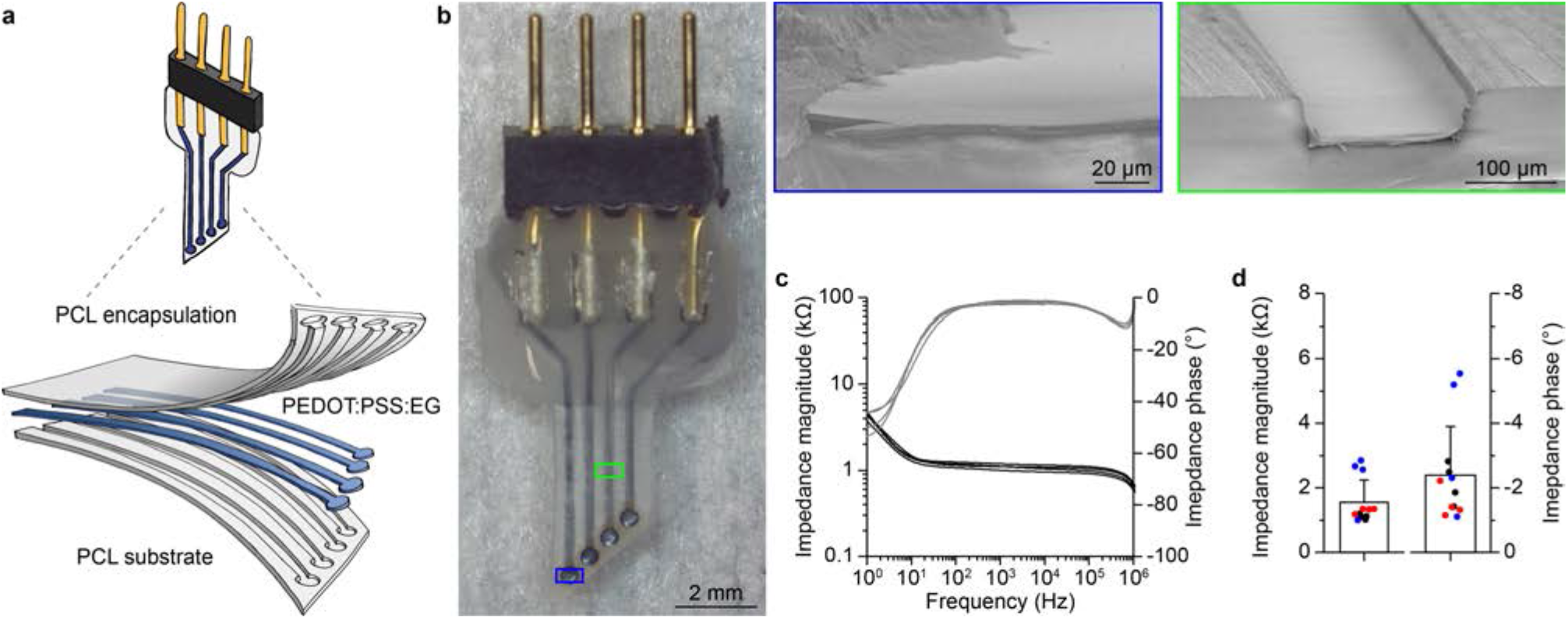
All-polymeric transient neural probes. **a**, Sketch of the TNP. **b**, Picture of a fully assembled TNP with four PEDOT:PSS:EG electrodes. Inserts show scanning electron microscope cross-sections of one electrode (image tilt 10 °) and one trace without encapsulation layer (image tilt 10 °). **c**, Plots of the impedance magnitude (black) and impedance phase (grey) of 4 electrodes from a representative TNP. **d**, Quantification (mean ± s.e.m.) of the impedance magnitude (left) and phase (right) at 1 kHz for all the 4 electrodes from 3 TNPs (respectively in red, blue, and black).

Electrical impedance spectroscopy (EIS) showed remarkable performances of all-polymeric TNPs based on PEDOT:PSS:EG, characterized by low impedance magnitude and low impedance phase (**Fig. 1c,d**). PEDOT:PSS:EG electrodes showed a low and stable impedance magnitude over a wide range of frequency (10 Ω to 1 MΩ), which makes them appropriate for the recording of both low- and high-frequency neuronal signals, such as local field potentials (LFPs) and neural spiking activities respectively. The comparison of multiple probes showed very low intra-probe variability. On the other hand, the inter-probe variability might be explained by the manual deposition process of PEDOT:PSS:EG resulting in a variable thickness of the conductive layer. To verify that the excellent electrochemical characteristics of the TNPs were not related to shortcuts between electrodes caused by the porosity of the PCL scaffold, we took advantage of an all-polymeric TNP with a modified design embedding 17 electrodes (**Supplementary Fig. 3**). In this specific device, four traces were interrupted, due to a gap in the PEDOT:PSS:EG layer (**Supplementary Fig. 3a-c**). As expected, only the corresponding electrodes were rightfully not functional (**Supplementary Fig. 3d,e**), thus confirming that shortcuts between electrodes caused by the porosity of PCL are not present. Cytotoxicity tests performed in-vitro showed a cell viability of 96.9 ± 7.2 % (mean ± s.d., 2 probes, 3 replicas per probe) and opened the way for the in-vivo validation of the devices (**Fig. 2**).

**Figure 2.**
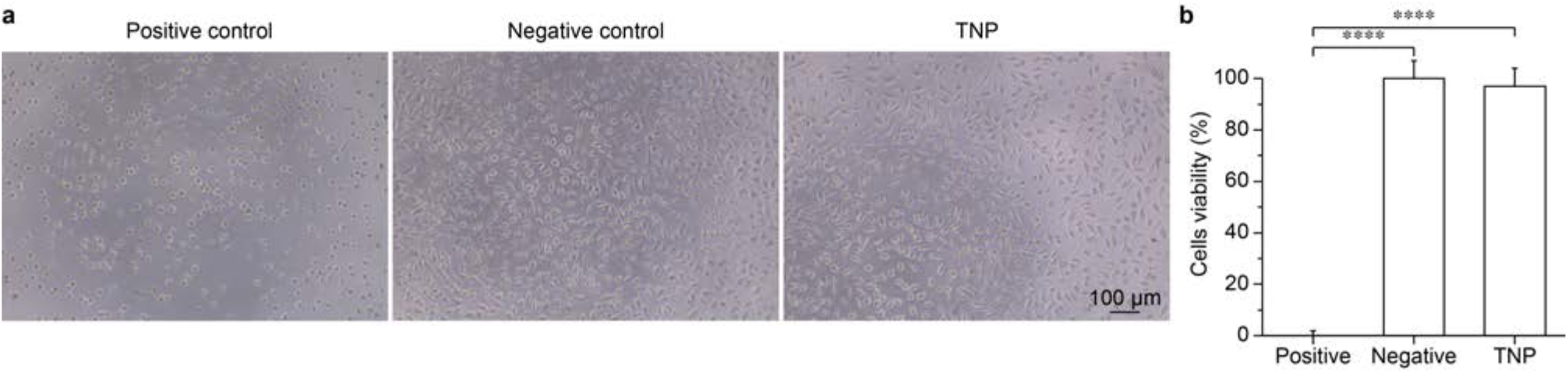
Cytotoxicity in-vitro. **a,** Representative optical images for each condition tested: positive control (left), negative control (middle), and TNP (right). **b**, Quantification of the mean (± s.d.) cells viability in the three conditions tested (3 replicas per condition). All-polymeric TNPs 96.9 ± 7.2 % (2 probes, 3 replicas per probe); positive control 0 ± 1.8 %; negative control 100 ± 6.8 %. One-way ANOVA: F = 853.4 and p < 0.0001. Tukey’s multiple comparisons test: positive control vs. negative control p < 0.0001; positive control vs. neural probes p < 0.0001; negative control vs. neural probes p = 0.4852.

### Functional validation in-vivo

To assess the capability of all-polymeric TNPs in recording neural activity, we performed in-vivo acute experiments in mice upon induction of epileptic seizures. All-polymeric TNPs were implanted in the visual cortex area of anaesthetised mice (**Fig. 3a**), and the neural activity was recorded both before and after intraperitoneal injection of the convulsant pentylenetetrazol (PTZ), which is routinely used to test anticonvulsants in animals[35–37]. The transition between resting state and epileptic activity after PTZ injection was clearly detected by the TNPs with an excellent signal to noise ratio (**Fig. 3b**), thus demonstrating the potential of these all-polymeric TNPs for monitoring epileptic activity.

**Figure 3.**
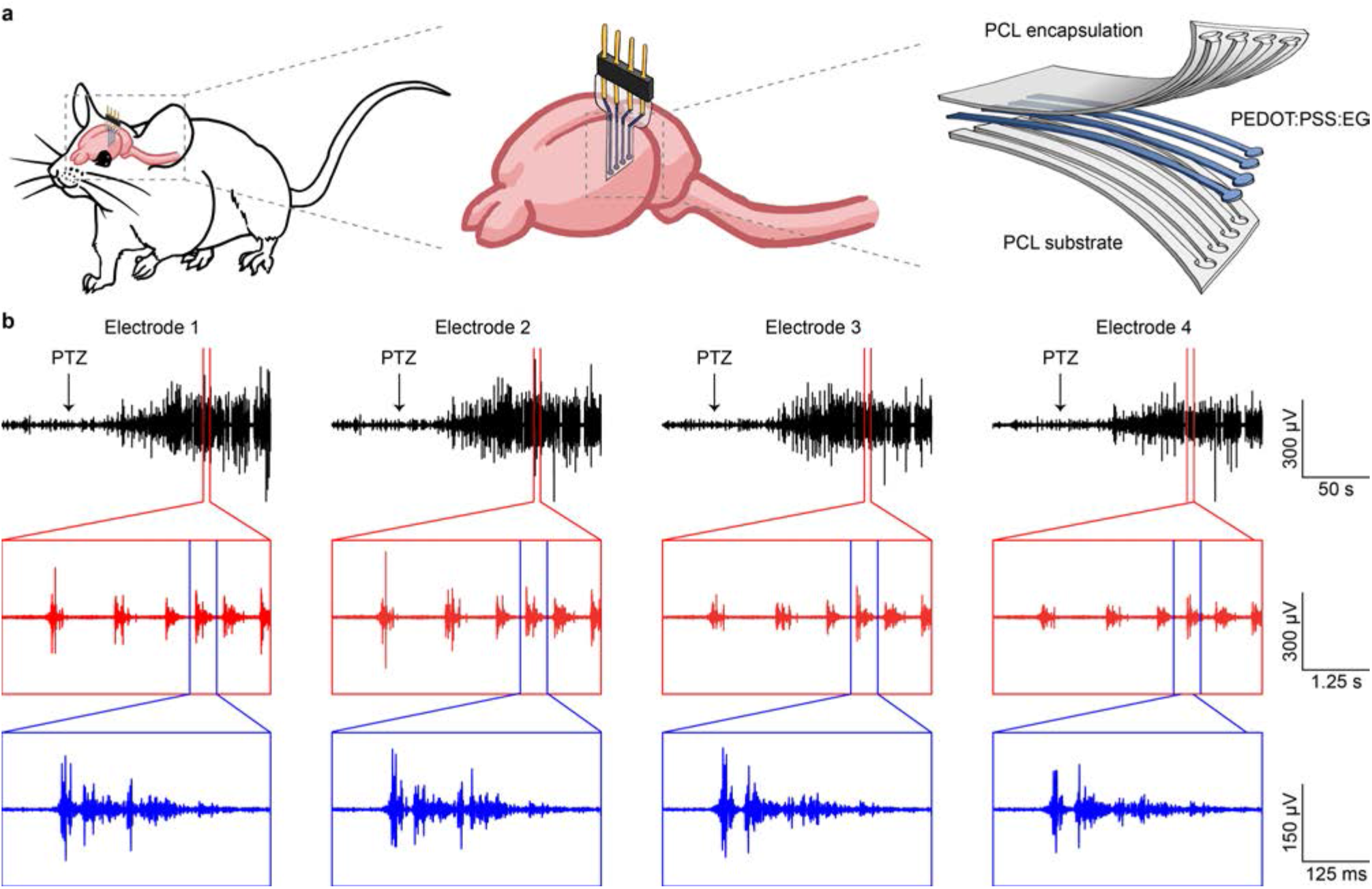
Acute recordings with all-polymeric transient neural probes. **a**, Sketch of the TNP implanted in a mouse brain for in-vivo recordings. **b**, Detection of epileptic activity after injection of PTZ (black arrow) from the four electrodes of a PEDOT:PSS:EG all-polymeric TNP. The red and blue boxes show an enlargement of the recorded activity.

Next, we performed chronic in-vivo recordings of baseline activity, LFPs at rest and visually evoked potentials (VEPs) to assess the longevity of the device. All-polymeric TNPs were implanted in the visual cortex area of mice, and their functionality was evaluated through periodic recordings up to 3 months post-implantation (**Fig. 4**). The broad-band noise of recordings was computed from baseline recordings (2.5 min, bandpass filter 0.1 - 2000 Hz and sampling frequency 8192 Hz). The noise plot (**Fig. 4a**) shows the evolution of the noise level for each working electrode over time (filled circles). Once an electrode was found not correctly working (open circles), it was excluded from further analysis. The applied criteria of exclusion were the following: i) appearance of periodic artefacts not linked to any biological function (red arrow in **Fig. 4a**); ii) noise level higher than the average plus twice the standard deviation of the noise at each time point; iii) lack of VEPs. The number of working electrodes is progressively reduced over time (**Fig. 4b**), with 3 out of 16 electrodes still active after three months. LFPs (2.5 min, bandpass filter 0.1 - 100 Hz and sampling frequency 819.2 Hz) are not strongly affected by broad-band noise because of the low pass filter at 100 Hz. On the other hand, LFPs are characterized by a strong delta rhythm (0.5 - 4 Hz) due to the animal anaesthesia (**Fig. 4c**). Early after implantation, electrodes showed a narrow power spectral density (PSD) in the range of delta rhythm (black traces in **Fig. 4c** and black lines in **Fig. 4d**). Over time, because of the electrode degradation, the normalized PSD shows activity at higher frequency in electrodes that were previously considered not working because of high broad-band noise (grey line in **Fig. 4d**). Conversely, electrodes still considered working showed a narrow PSD (red line in **Fig. 4d**). As a consequence, the relative power of the delta band decreases with the degradation of the electrodes (**Fig. 4e**). For each electrode, early after implantation, a low noise correlates with a high relative power of the delta band. However, at the exclusion point, there is a reduction of the relative delta power which is globally proportional to the increase of noise (linear regression: slope = −1.369, R square = 0.527). Overall, all-polymeric TNPs retain good in-vivo recording capabilities for months after implantation, even if they progressively lose functionality and only a few electrodes are still operational after three months.

**Figure 4.**
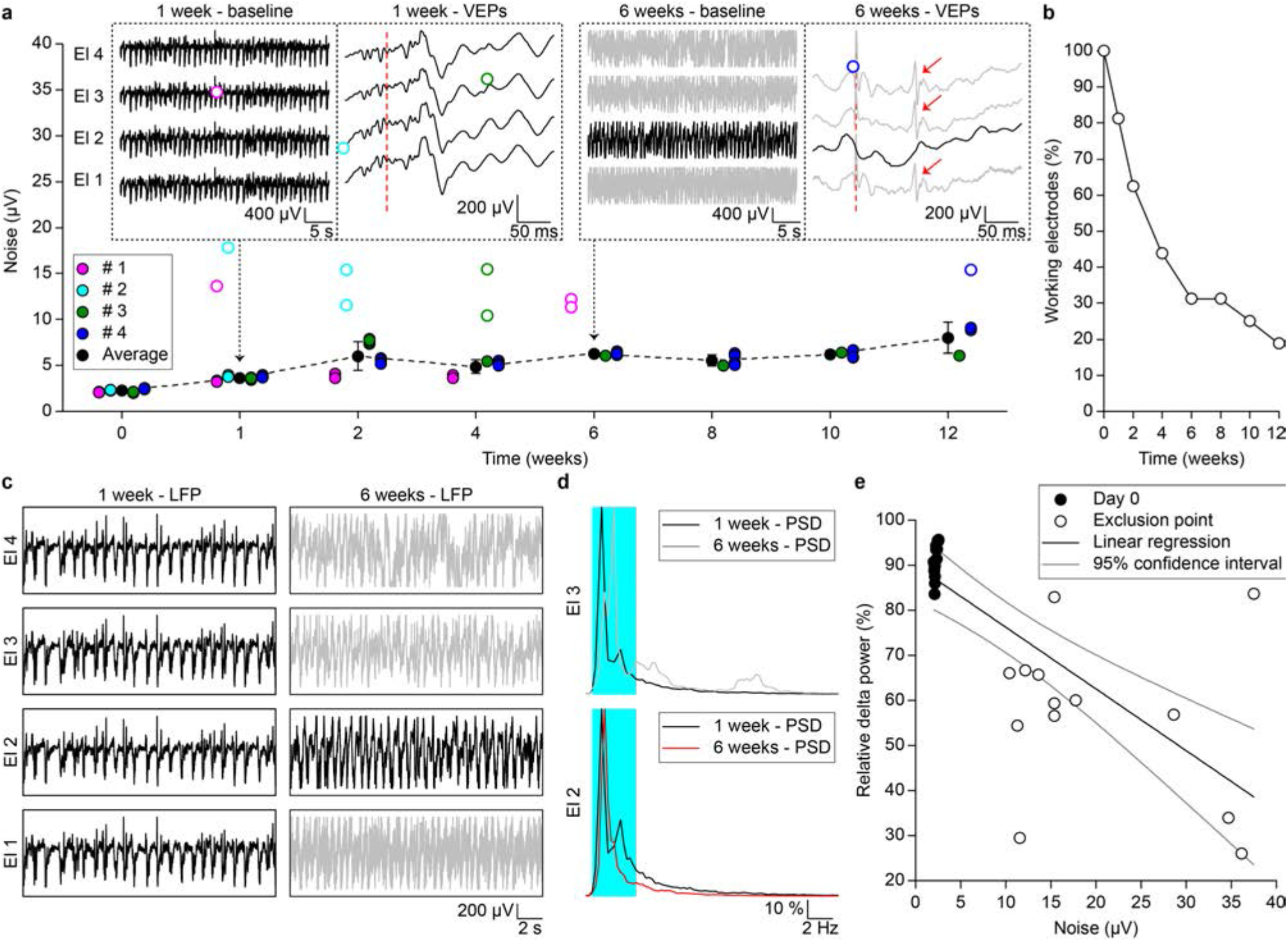
Long-term functioning in-vivo. **a**, Noise evaluation as a function of time during in-vivo chronic recordings at fixed time points. Different colours correspond to different mice; filled circles correspond to the electrodes considered functional and thus included in the calculation of the average (± s.d.) noise values (black circles and dashed line), while open circles correspond to the electrodes considered not correctly working and thus excluded from the average noise calculation. The two insets show single baseline recordings (left) and VEPs (right) from all the four electrodes of the TNP implanted in mouse # 3 at week 1 and week 6 postimplantation; the red dashed line indicates the light stimulus for VEPs (10 cd s m^-2^). The traces in grey are from electrodes considered not functional anymore. The red arrows show periodic artefacts not linked to any biological function. **b**, Plot of the percentage of working electrodes as a function of time during in-vivo chronic experiments. **c**, Representative example of LFPs recorded from all the electrodes of the TNP implanted in mouse # 3 at week 1 and week 6 post-implantation. The traces in grey are from electrodes considered not functional anymore. **d**, Representative examples of normalized power spectral densities obtained from the recordings from two electrodes (numbers 2 and 3) of the probe implanted in mouse # 3 at week 1 (black traces) and week 6 (black grey and red) after implantation. The trace in grey is from the electrode considered not functional anymore, while the trace in red is from the electrode considered still functional. **e**, Correlation between the relative power of the delta band and the noise of the electrodes immediately after implantation (16 electrodes, black circles) or at the time point of exclusion (13 electrodes, open circles). The black line is the linear regression, while the grey lines the 95% confidence interval.

### Evaluation of the foreign body reaction

To investigate the acute phase of the foreign body reaction induced by TNPs, we performed a histological paired comparison (1- and 2-months post-implantation, 6 mice for the first time point and 5 mice for the second time point) between all-polymeric TNPs and polyimide (PI) implants of similar dimensions (thickness 125 μm), surgically placed in the two hemispheres of the brain within the same mouse (**Fig. 5a**). The PI implants were chosen as a reference for non-degradable flexible implants. Also, the implants were made with similar dimensions to be more comparable. In this analysis, for each probe, horizontal brain slices sampled at different cortical depths (from 1 to 4) were analysed and averaged. Also, for each cortical depth, three slices were averaged. The staining for the glial fibrillary acidic protein (GFAP) (**Fig. 5b,c**) and the corresponding quantification analysis (**Fig. 5d**) performed on segmented images (**Supplementary Fig. 4**) showed that the area occupied by the GFAP signal in the zone adjacent to the probe (as a percentage of total area, **Supplementary Fig. 4d**) exhibits a decreasing trend over time for all-polymeric TNPs (p = 0.0178; two-tailed unpaired t-test) and a stable trend for PI probes (p = 0.6924; two-tailed unpaired t-test). Also, the staining for the cluster of differentiation 68 protein (CD68), a marker of phagocytosis in microglia[38], and the corresponding quantification analysis (**Fig. 5b,c,e**) showed that the area occupied by the CD68 signal exhibits a slightly increasing trend over time for all-polymeric TNPs (p = 0.2897; two-tailed unpaired t-test), while it exhibits a decreasing trend over time for PI probes (p = 0.5905; two-tailed unpaired t-test). Interestingly, a different analysis of the segmented images (**Supplementary Fig. 5**) performed at the level of the electrode (slice 4 in **Fig. 5b,c**), the only point where the PEDOT:PSS:EG is exposed to the tissue, showed that the CD68 signal is highly clustered on the side of the PEDOT:PSS:EG electrodes at both time points (**Fig. 5f,g**). The probes were always implanted with the PEDOT:PSS:EG layer exposed to the caudal part of the brain to discriminate the two sides during image analysis.

**Figure 5.**
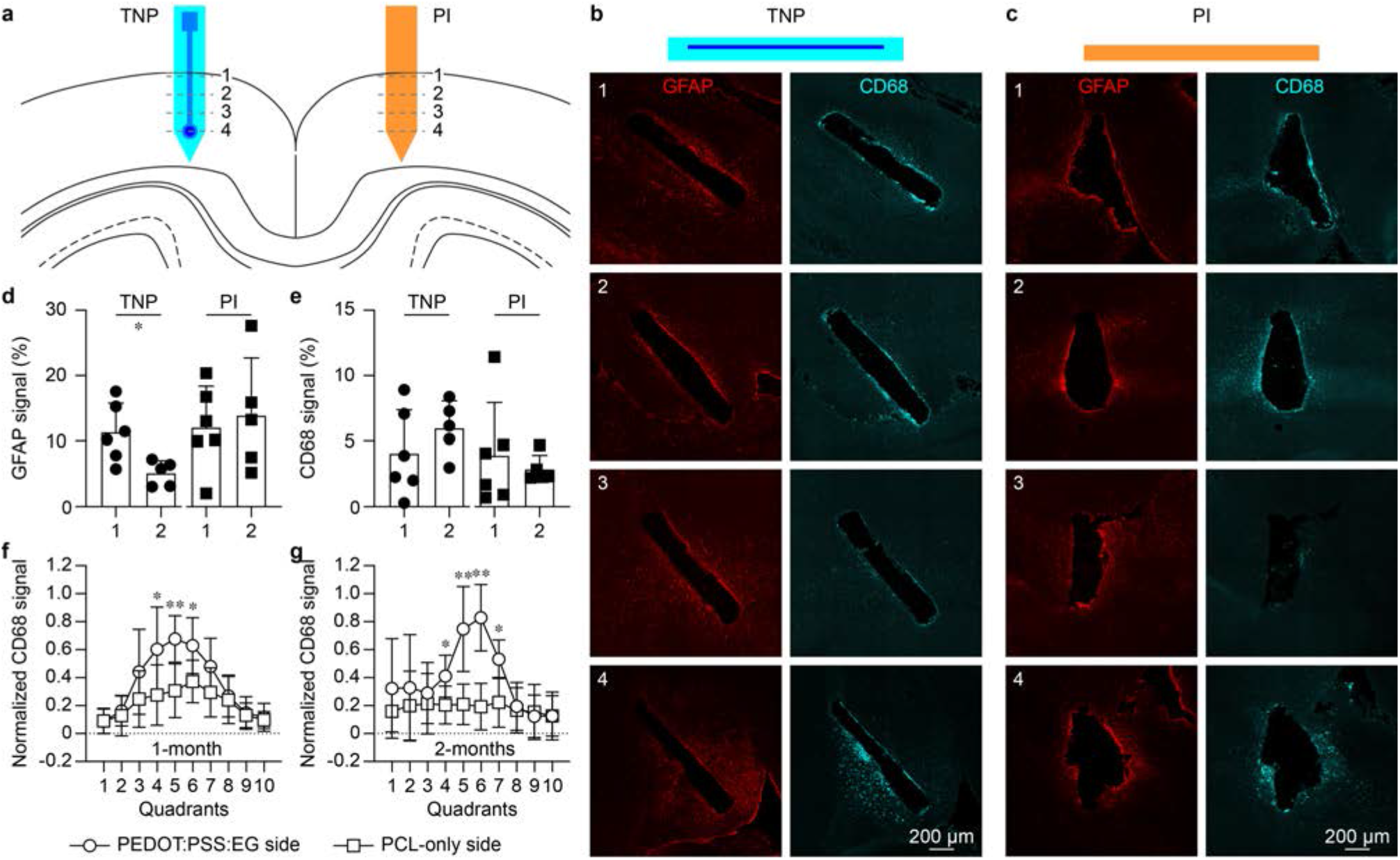
Foreign body reaction induced by all-polymeric transient neural probes. **a**, Sketch of the TNPs and PI probes implanted in both visual cortices of mice. For each probe, four slices were taken at four depths into the cortical layers. **b,c**, Example of GFAP (in red) and CD68 (in cyan) expression in horizontal slices (four depths into the cortical layers indicated by the numbers) obtained from the hemisphere implanted with the TNP (**b**) and with the PI probe (**c**) 1-month post-implantation. **d,e**, Quantification of the area occupied by GFAP signal (**d**) and CD68 signal (**e**) cells at 1-month and 2-month post-implantation for TNPs and PI probes (results from all the slides at each depth were averaged together). **f,g**, Quantification of the normalized area occupied by the CD68 signal cells above (PEDOT:PSS:EG side) and below (PCL-only side) TNPs at the level of the electrode at 1-month and 2-month post-implantation (* indicates p < 0.05 and ** indicates p < 0.01; two-tailed unpaired t-test).

This evidence suggested the presence of PEDOT:PSS:EG dethatched from the PCL substrate stimulates phagocytic activity of microglial cells. To further investigate this hypothesis, we qualitatively assessed the interaction between phagocytising microglial cells and PEDOT:PSS:EG. To this end, we fabricated a TNPs with a PCL substrate coated with PEDOT:PSS:EG labelled with the fluorescent marker fluorescein isothiocyanate (FITC)[19] and implanted them into the cortex of mice expressing tdTomato in microglia (**Fig. 6a**). In this experiment, to accelerate the delamination of PEDOT:PSS:EG from PCL, all-polymeric TNPs were not encapsulated. High-resolution imaging with a clarity optimized light-sheet microscope (COLM) on clarified brains 1-month (**Fig. 6b**) and 2-months (**Fig. 6c**) post-implantation showed colocalization of PEDOT:PSS:EG (in green) and microglial cells (in red), suggesting that the microglia is participating in the clearance of PEDOT:PSS:EG from the implantation site. Horizontal brain sections at 1-month postimplantation also revealed microglial cells (in red) localizing in the region were PEDOT:PSS:EG (in green) is delaminated or directly in contact with the brain (**Fig. 6d**). These cells are CD68 positive (in white), indicating phagocytic activity. Although the precise extension of the glial scar is difficult to quantify in brain slices because of the high variability within samples (**Fig. 5**), based on the results discussed above, we hypothesise that implanted TNPs lead to the formation of a less tight glial scar compared to PI probes. As a consequence, the microglia has the space needed to make its way to the probe and phagocyte the delaminated PEDOT:PSS:EG flakes.

**Figure 6.**
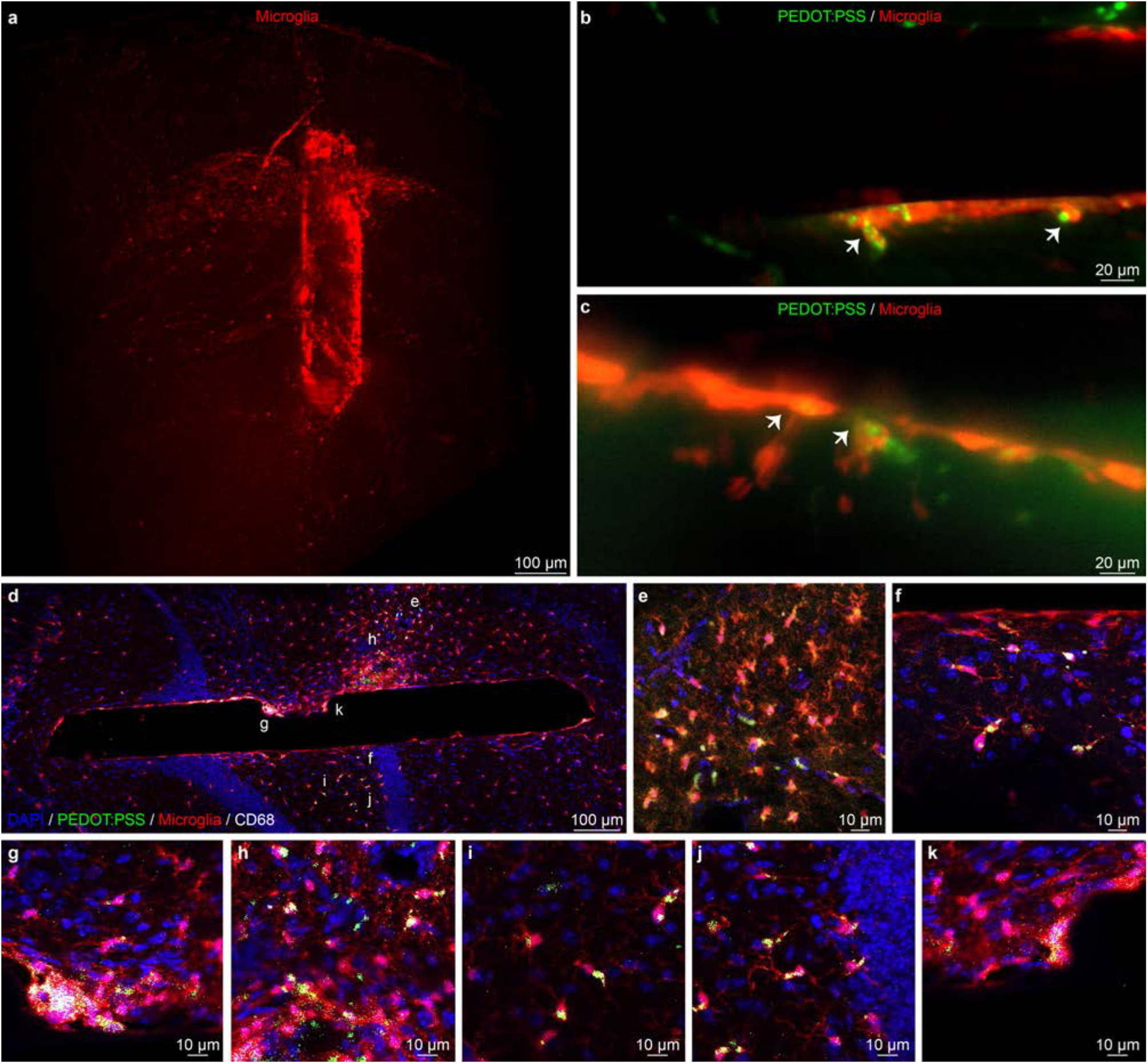
Integration with microglia. **a**, 3D mesoSPIM image of a clarified brain sample with fluorescent microglia (red) implanted with a TNP with FITC-labelled PEDOT:PSS at 1-month post-implantation. **b,c**, COLM images of the probe surface, showing fluorescent PEDOT:PSS (in green) and microglia (in red) colocalization 1-month (**b**) and 2-months (**c**) post-implantation. Microglia engulf PEDOT:PSS:EG particles which are detaching from the probe (white arrows). **d**, Horizontal brain section of mouse brain with fluorescent microglia (in red) implanted with a fluorescent TNP (without encapsulation) at 1-month post-implantation. **e,k**, Various magnifications of the section in **d** (corresponding letters) showing phagocytic microglia engulfing PEDOT:PSS:EG (in green). The expression of CD68 (in white) is detected in phagocytic cells.

### Probe’s degradation

One of the most exciting features of transient medical devices is their natural ability to disappear in the body, which in turns reduce the long-term foreign body reaction. PCL bulk degradation occurs slowly with a time scale spanning several months or years[23], thus making an in-vivo assessment in animals extremely challenging. In order to investigate the degradation process of all-polymeric TNPs, we performed in-vitro accelerated degradation tests, by leaving 3 TNPs immersed in phosphate-buffered saline at 37 °C and pH 12 and monitoring their weight over time. Scanning electron microscope images showed the appearance of micro- and macro-cracks over time due to hydrolysis (**Fig. 7a**), and all-polymeric TNPs undergoes full degradation in about one year of time at pH 12 (**Fig. 7b**). A comparison between the days to reach a weight loss of 5 % at pH 12 (42 days) and at pH 7.4 (325 days) revealed that our test has an acceleration factor of about 7.7, which is in line with previous reports for PCL (8.5)[26]. Slow degradation could be an appealing feature for TNPs, since it might allow for better tissue remodelling and reduced chronic trauma at the implantation site[26,39,40]. To provide evidence of degradation in-vivo of all-polymeric TNPs, we implanted them in the cortex of mice and subsequently performed immunofluorescence assays on horizontal brain slices at a fixed time point. Upon long-term implantation (9 months), the all-polymeric TNPs showed a minor chronic glial scar (**Fig. 7c**), evaluated with staining for astrocytes against GFAP. The formation of a glial scar should be minimized or avoided for several reasons: it increases the distance between the recording surface of the electrode and neurons, eventually impairing the recording capabilities of the device[41,42], and acts as a barrier for growth factors, ions, and signalling molecules, thus promoting neural apoptosis and impeding axon regeneration[43,44]. In the case of TNPs, the presence of a limited glial scar allows tissue remodelling around the probe, and, as a consequence, neurons (labelled against NeuN in green) are free to migrate into the device, which serves as a support scaffold during the degradation (**Fig. 7d**). This evidence is a proof of the probe’s degradation in-vivo and a promising sign of reduced chronic trauma.

**Figure 7.**
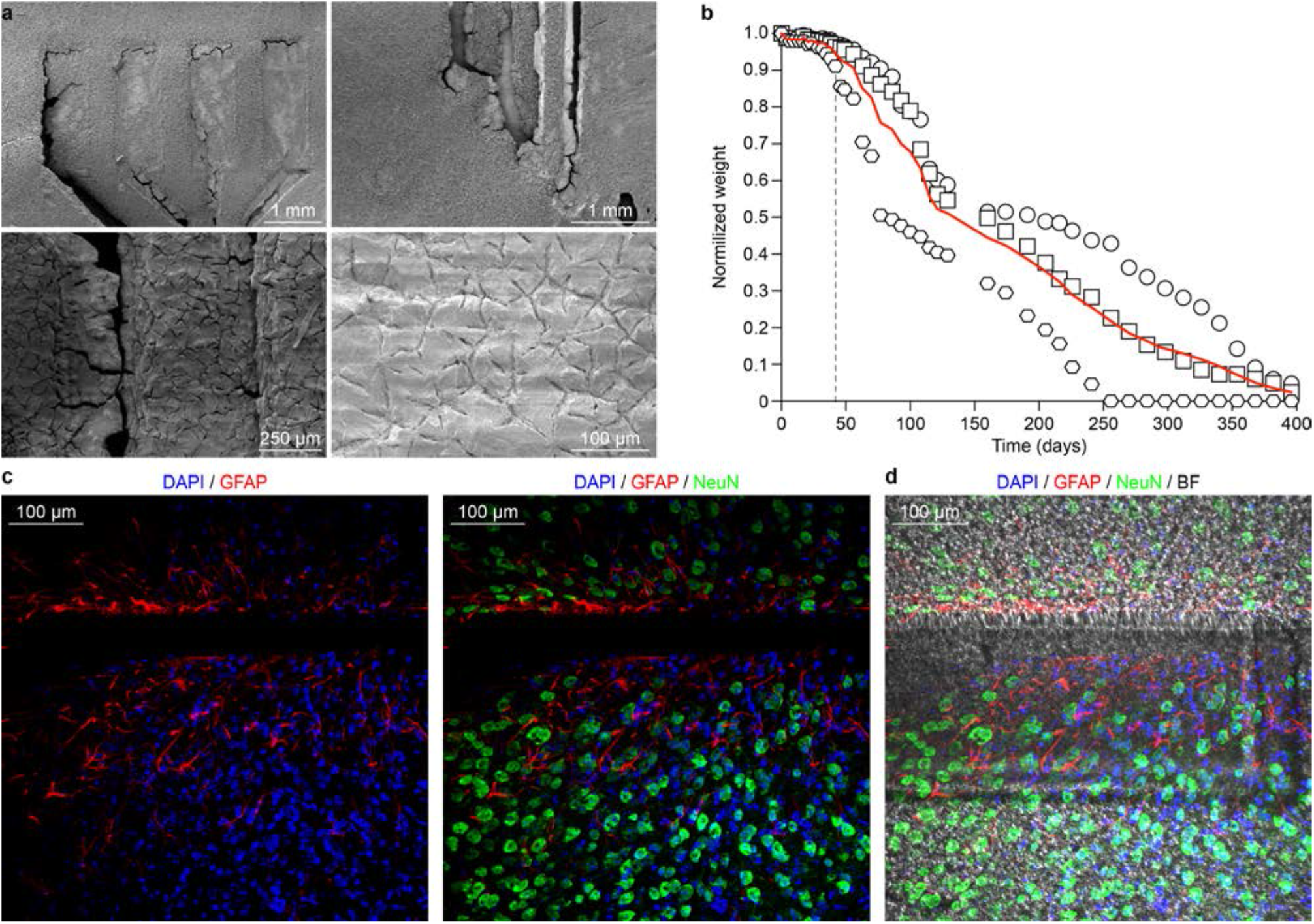
Degradation of all-polymeric transient neural probes. **a**, Scanning electron microscope images at several magnifications of one all-polymeric TNP obtained 191 days after soaking in phosphate-buffered saline solution at 37 °C and pH 12. **b**, Plot of the normalized probe weight versus time of 3 TNPs during a degradation test in saline solution at 37 °C and pH 12. The red line is the average of the 3 TNPs. The grey dashed line is the time point (day 42) corresponding to a 5 % weight loss. **c**, Horizontal brain section from a mouse implanted with a TNP in the cortex. 9-months post-implantation staining against GFAP and NeuN. **d**, The same image in **c** with a bright-field overlay to highlight the area occupied by the TNP.

## Discussion

In biomedical engineering and medicine, transient devices are relatively a new concept to widen the application possibilities of implantable medical devices[6]. From our perspective, it was worth exploring this new concept with fully polymer-based devices to exploit their unique characteristics and try to overcome some of the current limitations. So far, transient medical devices showed a short functional lifetime because of the fast degradation of transient metal in contact with body fluids[11,18].

In this work, we showed the fabrication and characterization of a polymer-based implantable device conceived to disappear after being functional for a few months rather than a few days. A longer device lifetime will allow, for instance, the development of prosthetic devices for mid-term monitoring of the brain activity either before or after surgery in chronic epileptic patients. Nowadays, clinicians make decisions on localized surgical treatments based on intracranial electroencephalography recordings that are typically lasting for only 1 or 2 weeks in favour of patient compliance. On the other hand, recent publications in this field revealed that intracranial electroencephalography recordings do not stabilize until several weeks after the device’s implantation, due to the invasiveness of the procedure and the inflammatory response of the tissue[45,46]. Therefore, an extended chronic monitoring period, together with the possibility to mitigate the inflammatory reaction, would enable a more accurate representation of the spatio-temporal variability of the seizures, translating into a better diagnosis for the patient. Furthermore, a transient device with extended recording capability would allow for continuous monitoring after the surgical procedure, thus providing essential insights into the brain activity during the recovery period and possibly on the effects of the following pharmaceutical treatment. Also, the total absence of metals in the device would provide magnetic resonance imaging compatibility[29,47,48], allowing the pairing of the electrophysiological monitoring with structural and functional images, therefore obtaining a clearer picture of the brain activity during the recovery period.

Even though only a few electrodes (approximately 20%) were still operational three months post-implantation, we believe that there is still room for fabrication strategies that can improve the device lifetime, such as enhancing the adhesion between PEDOT:PSS and PCL. Also, even if a performance comparison with other inorganic and not-transient devices is out of the scope of this study (due to the transient nature of the presented device), the electrochemical characterization of all-polymeric TNPs showed a stable performance within a wide range of frequencies, thus allowing us to perform acute and chronic in-vivo recordings in mice with excellent signal to noise ratio.

A deep study of the foreign body reaction revealed that all-polymeric TNPs were responsible for a minimal glial scar around the implant. As a consequence, neurons were able to colonize the slowly degrading PCL layers, hence promoting tissue remodelling: such behaviour is the opposite to what is usually observed with permanent implants, where neurons are depleted from the implantation site[41].

In summary, the in-vivo validation and the degradation study, together with preliminary evidence of phagocytic microglia engulfing the conductive polymer demonstrate that the combination of the transient electronic approach and the usage of polymers is a new promising path to improve the device-tissue interface, extend the durability of implanted transient devices, and increase their possible applications. Although further research is needed to fully understand the fate of microglial cells upon phagocytosis of PEDOT:PSS:EG, these results can drive the conceptualization of new conductive polymers with links that are degradable for example by proteolytic enzymes present in resident microglia.

## Acknowledgement

This work has been supported by École polytechnique fédérale de Lausanne, Medtronic and European Commission (EU project 701632). The authors would like to thank Dr. Laura Batti and Audrey Tissot for the support with the brain clarification process and imaging and Vivien Gaillet for the drawings.

## Author contributions

L.F. designed the study, fabricated the devices, performed electrochemical characterization and analysed data, performed surgeries, performed immunohistochemistry, collected confocal images, and wrote the manuscript. P.V. designed the study, performed surgeries, performed electrophysiology and analysed data, performed immunohistochemistry, collected and analysed confocal images, and wrote the manuscript. E.G.Z. performed the surgeries and immunohistochemistry, collected confocal images, and performed cytotoxicity assay. A.F. fabricated the devices and performed the accelerated degradation study in-vitro. K.M. and R.C.P. supplied genetically modified mice and participated in the histological tests. D.G. supervised the entire study and wrote the manuscript. All the authors read and accepted the manuscript.

## Supplementary information

**Supplementary Figure 1.**
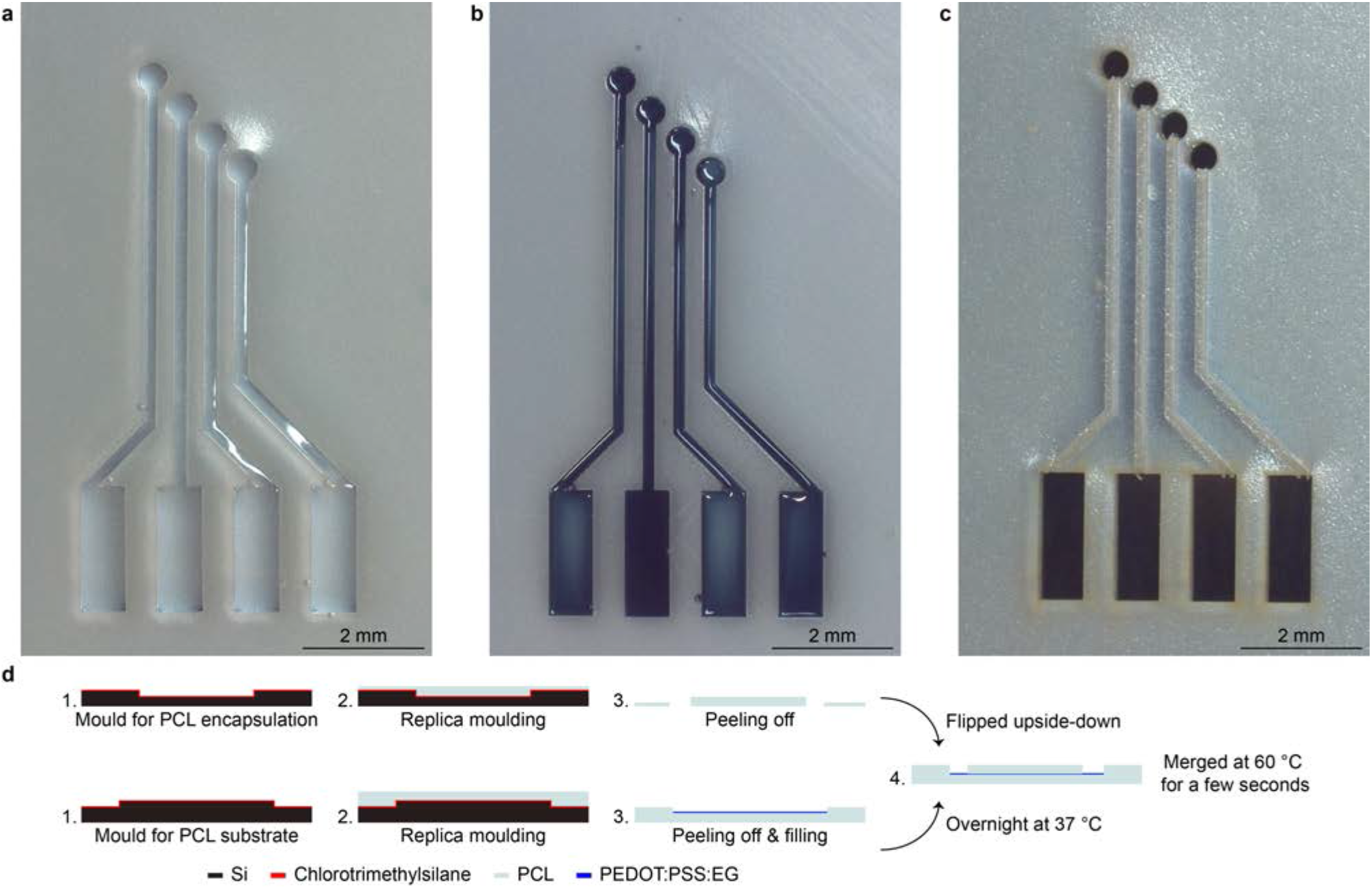
Fabrication of all-polymeric transient neural probes. **a**, PCL substrate obtained via replica moulding. **b**, PCL substrate with PEDOT:PSS:EG deposited in the hollow regions (50-μm depth) corresponding to the electrodes, traces, and pads. **c**, Probe encapsulated with a matching PCL layer with protruding traces (60-μm thick) and openings to expose the recording sites and the pads. **d**, Process flow of the main fabrication steps (dimensions are not in scale).

**Supplementary Figure 2.**
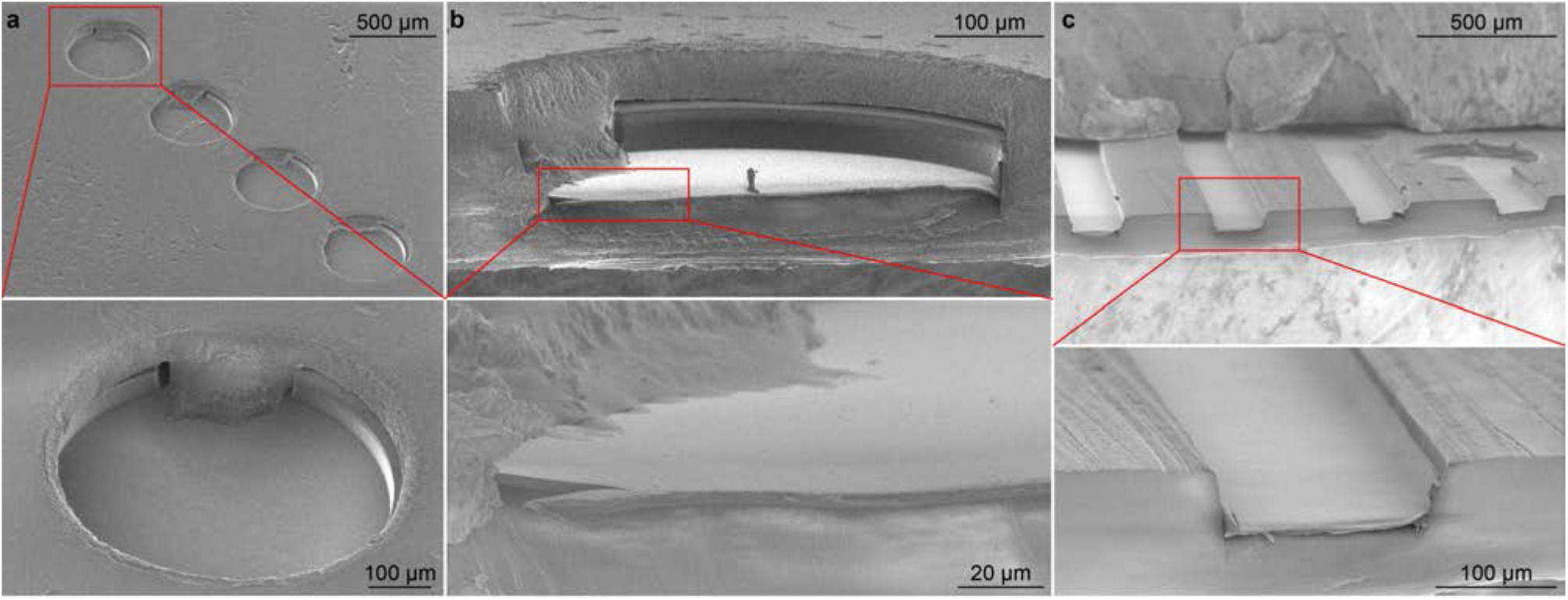
Scanning electron microscope characterization of an all-polymeric transient neural probe. **a**, Image of the four electrodes encapsulated in PCL (image tilt 45 °). The insert shows a magnification of one electrode (image tilted 45 °). **b**, Cross-section of one electrode after cutting (image tilt 10 °). The insert shows a magnification of one electrode (image tilted 10 °). **c**, Cross-section at the level of the traces after cutting without the encapsulation layer (image tilted 10 °). The insert shows a magnification of one electrode (image tilted 10 °).

**Supplementary Figure 3.**
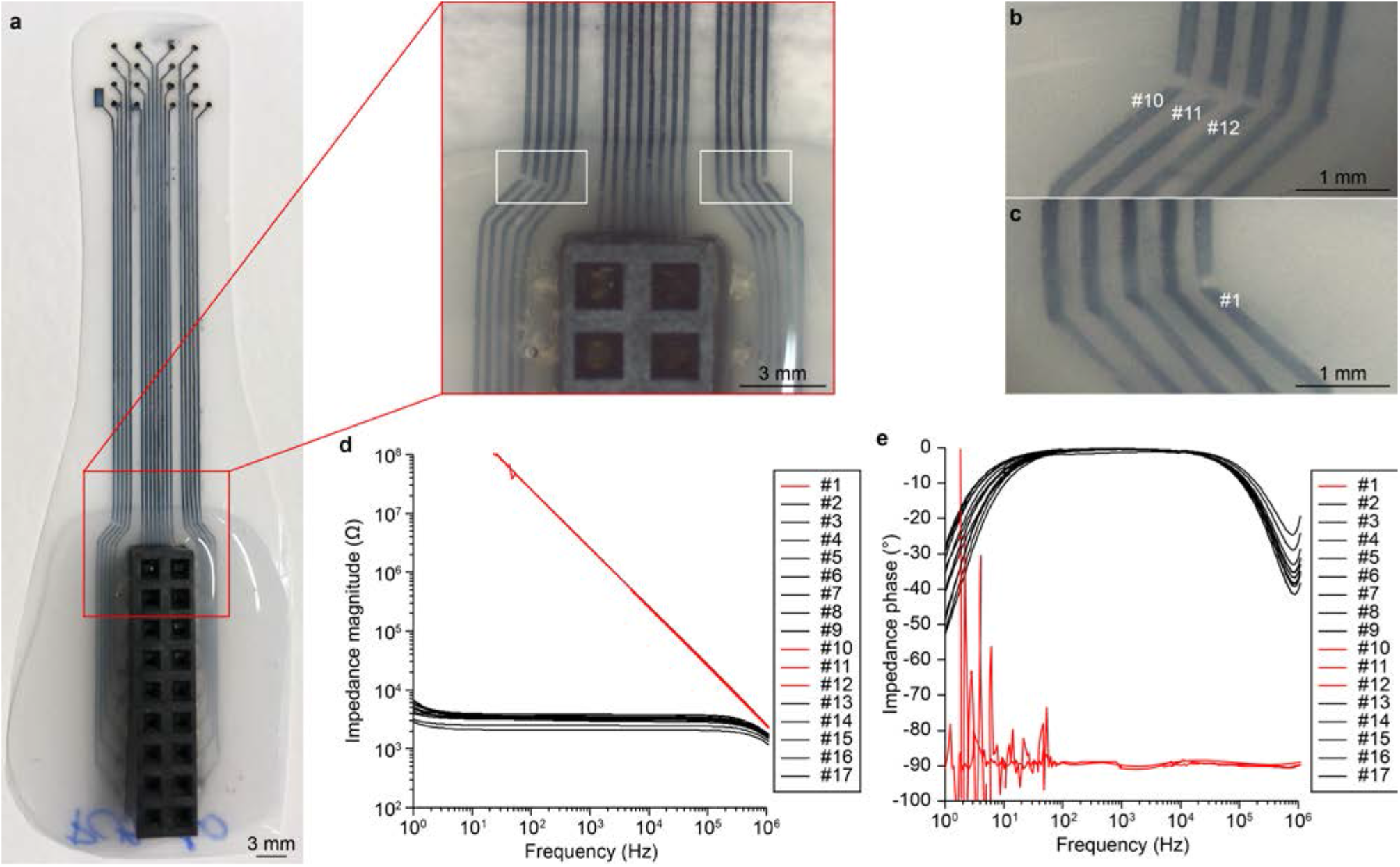
Independence of the electrodes. **a**, Picture of an all-polymeric TNP with higher channel count (17 electrodes, 700-μm in diameter) and its connector. The red box shows a magnified view of a portion of the neural probe where traces appears to be disconnected. **b,c**, Magnification of the disconnected traces and corresponding electrode numbers. The images correspond to the white boxes in **a. d**, Plot of the impedance magnitude of all 17 electrodes as a function of frequency. In red are highlighted the measurements from the electrodes with disconnected traces. **e**, Plot of the impedance phase of all 17 electrodes as a function of frequency. In red are highlighted the measurements from the electrodes with disconnected traces.

**Supplementary Figure 4.**
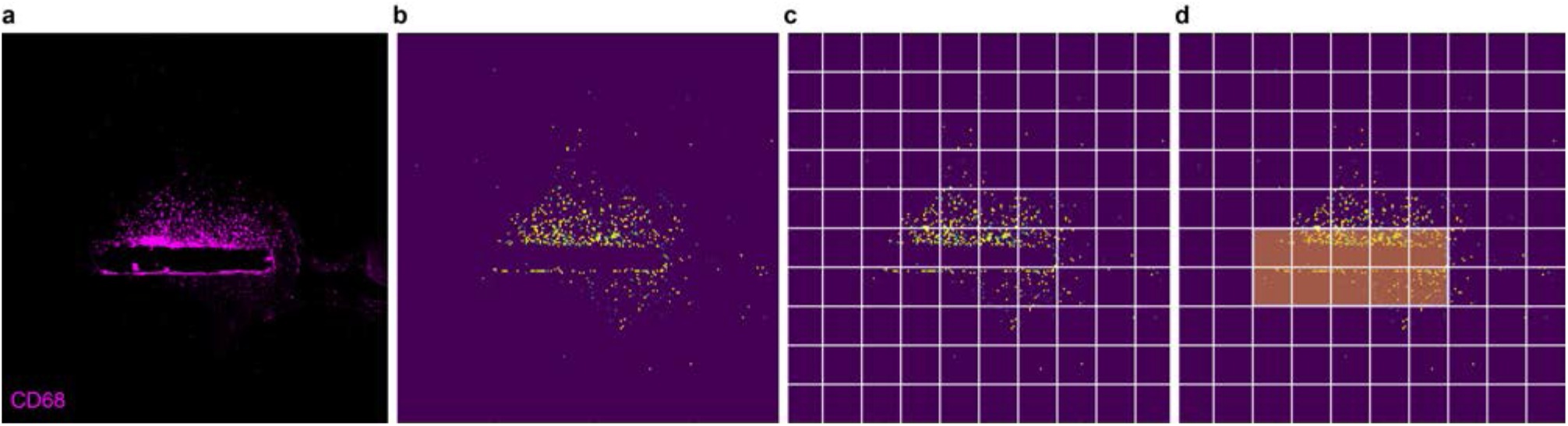
Image segmentation. **a**, Representative image from a horizontal section of the mice brain implanted with an all-polymeric TNP and stained against CD68. **b**, Binary image obtained using a local threshold algorithm on the image in **a**. To the pixels whose intensity is above the threshold it is assigned the value 1 (yellow), while to the other pixels it is assigned the value 0 (background). **c**, The image in **b** is divided into 10 x 10 quadrants, and the total area of the blobs (formed by adjacent pixels with value 1) is computed for each quadrant. **d**, The proximal area (area of the zone adjacent to the probe) is defined as the cumulative area of all the quadrants that are included in a rectangle defined by the extremities of the probe minus the area of the probe itself.

**Supplementary Figure 5.**
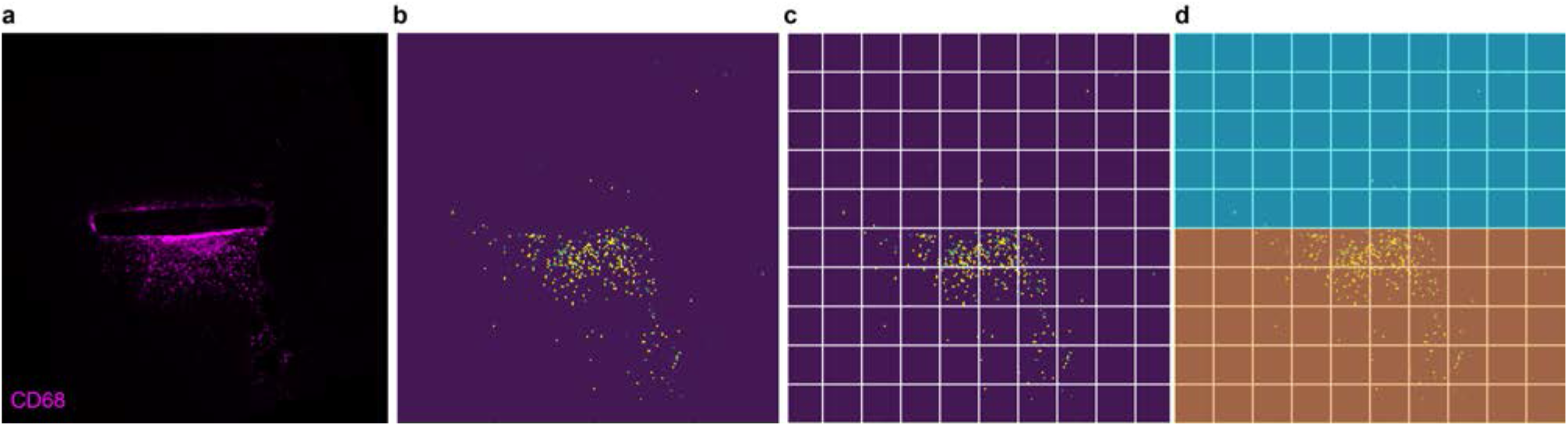
Differential image segmentation above and below the probe. **a**, Representative image from a horizontal section at the electrode level of the mice brain implanted with an all-polymeric TNP and stained against CD68. **b**, Binary image obtained using a local threshold algorithm on the image in **a**. To the pixels whose intensity is above the threshold it is assigned the value 1 (yellow), while to the other pixels it is assigned the value 0 (background). **c**, The image in **b** is divided into 10 x 10 quadrants, and the total area of the blobs (formed by adjacent pixels with value 1) is computed for each quadrant. **d**, The image is divided into two parts: the side of the probe with PEDOT:PSS:EG exposed (yellow) and the side of the probe with only the PCL exposed (blue).

